# Fungal adaptation to plant defenses through convergent assembly of metabolic modules

**DOI:** 10.1101/343673

**Authors:** Emile Gluck-Thaler, Vinod Vijayakumar, Jason C. Slot

## Abstract

The ongoing diversification of plant defense compounds exerts dynamic selection pressures on the microorganisms that colonize plant tissues. Evolutionary processes that generate resistance towards these compounds increase microbial fitness by giving access to plant resources and increasing pathogen virulence. These processes entail sequence-based mechanisms that result in adaptive gene functions, and combinatorial mechanisms that result in novel syntheses of existing gene functions. However, the priority and interactions among these processes in adaptive resistance remains poorly understood. Using a combination of molecular genetic and computational approaches, we investigated the contributions of sequence-based and combinatorial processes to the evolution of fungal metabolic gene clusters encoding stilbene cleavage oxygenases (SCOs), which catalyze the degradation of biphenolic plant defense compounds known as stilbenes into monophenolic molecules. We present phylogenetic evidence of convergent assembly among three distinct types of SCO gene clusters containing alternate combinations of phenolic catabolism. Multiple evolutionary transitions between different cluster types suggest recurrent selection for distinct gene assemblages. By comparison, we found that the substrate specificities of heterologously expressed SCO enzymes encoded in different clusters types were all limited to stilbenes and related molecules with a 4’-OH group, and differed modestly in substrate range and activity under the experimental conditions. Together, this work suggests a primary role for genome structural rearrangement, and the importance of enzyme modularity, in promoting fungal metabolic adaptation to plant defense chemistry.

## Introduction

Metabolic pathways for the synthesis and degradation of secondary/specialized metabolites (SMs) govern critical interactions between organisms (Raguso *et al.* 2015). For example, plants produce numerous SMs that defend against fungal attack and decay, while fungi escape the inhibitory effects of these molecules by neutralizing or breaking them down (Michielse *et al.* 2012; Hammerbacher *et al.* 2013; Kettle *et al.* 2015). The co-diversification of such biosynthetic and degradative pathways may be a driving force behind global biodiversity and ecosystem function (Bailey *et al.* 2009; Futuyma & Agrawal 2009; Speed *et al.* 2015), affecting the spatial and temporal assembly of complex communities (Züst *et al.* 2012; Richards *et al.* 2015).

Many biosynthetic and degradative SM pathways in bacteria, plants and fungi are encoded in metabolic gene clusters (MGCs), which consist of neighboring sets of genes encoding enzymes, regulators and transporters functioning in the same metabolic processes (Nützmann *et al.* 2016; Adamek *et al.* 2017; Slot 2017). MGCs are more frequently affected by gene duplication and HGT compared with unclustered genes, and are hotspots for metabolic diversification through both combinatorial and sequence-based mechanisms (Wisecaver *et al.* 2014; Lind *et al.* 2017). Combinatorial-based diversification of biosynthetic and degradative pathways typically results from processes such as horizontal gene transfer (HGT) and/or the rewiring of regulatory circuits that assemble existing enzymatic reactions into qualitatively new pathways (Wagner 2011). Sequence-based diversification results from point mutations, insertions, deletions and recombination within enzyme-encoding genes that lead to specialized or novel enzymatic functions (Copley 2009). Changes to the coding sequences of individual genes within MGCs can contribute quantitatively (Lendenmann *et al.* 2014) and qualitatively (Proctor *et al.* 2008) to biochemical phenotypes, while changes to the combinations of genes within MGCs are often associated with the addition or deletion of particular enzymatic reactions from a given pathway (Yabe & Nakajima 2004; Ehrlich *et al.* 2004; Proctor *et al.* 2009). Although combinatorial and sequence-based processes are not mutually exclusive, their effects are rarely examined in parallel, precluding an assessment of their relative contributions to the diversification of adaptive metabolic pathways encoded in MGCs.

Among eukaryotes, fungi possess some of the most highly clustered genomes (Slot 2017). Fungal MGCs that synthesize SMs are well studied, and increasingly, fungal mechanisms to degrade/assimilate plant SMs are also found encoded in clusters (Kettle *et al.* 2015; Glenn *et al.* 2016; Gluck-Thaler & Slot 2018). For example, a fungal MGC encoding a putative pathway for the degradation of plant-produced stilbene molecules was recently reported by (Greene *et al.* 2014). Stilbenes are a structurally diverse group of ethene-linked biphenolic SMs produced by distantly related plant families and some bacteria (Chong *et al.* 2009). Stilbene biosynthesis has convergently evolved at least four times across the plant kingdom (Tropf *et al.* 1994; Han *et al.* 2014), and numerous plant lineages produce distinct suites of stilbene derivatives (Chong *et al.* 2009). Stilbenes are important components of constitutive and inducible plant defenses, often with broad-spectrum antifungal activity (Celimene *et al.* 1999; 2001; Jeandet *et al.* 2010), and some form lignin monomers that contribute to plant cell wall integrity (Del Río *et al.* 2017). The ability to degrade or neutralize stilbenes in some fungal pathogens can be directly associated with their ability to colonize host tissue (Hammerbacher *et al.* 2013; Wadke *et al.* 2016).

One stilbene catabolism mechanism entails the enzymatic cleavage of the 1,2-diphenylethylene stilbene backbone into two phenolic aldehydes by stilbene cleavage oxygenases (SCO; EC: 1.13.11.43) derived from the carotenoid cleavage dioxygenase family (Harrison and Bugg 2014). Five fungal SCOs from *Ustilago maydis* (*Um*Rco1; XP_761231), *Aspergillus fumigatus* (*Af*Rco1; XP_746307), *Chaetomium globosum* (*Cg*Rco1; XP_001219451), *Botryotinia fuckeliana* (*Bf*Rco1; XP_001548426) and *Neurospora crassa* (Cao-1; XP_961764) have been functionally characterized to date and were reported to share identical specificities towards multiple substrates (Brefort *et al.* 2011; Díaz-Sánchez *et al.* 2013). However, how the SCO-containing catabolic pathways adapt to diverse stilbene structures remains an open question.

Here, we investigated the evolution of fungal SCO function and the composition of associated metabolic gene clusters (*sco* clusters). We first identified three *sco* clusters with distinct gene combinations. Given that alternative types of MGCs typically encode different metabolic pathways, we predicted that SCOs encoded in different *sco* cluster types would either have 1) specificity to alternative substrates, suggesting enzyme functional divergence has priority over the formation of new cluster types, or 2) conserved specificity, suggesting cluster/pathway remodeling can operate independently of sequence-based evolution. We analyzed the substrate specificities of four heterologously expressed SCOs from different cluster types and unclustered genomic regions, and found evidence for broad conservation of substrate specificity in both unclustered and clustered SCOs, with only modest variation in the substrate specificity of SCOs found in alternate cluster types. Conversely, we found evidence for multiple independent origins of and repeated transitions between different types of *sco* clusters, consistent with selection acting upon recurrent combinations of enzyme-encoding genes. These results suggest that the reorganization of *sco* clusters may underpin adaptive shifts in stilbene degradation pathways, while changes in enzymatic function may play a secondary role.

## Methods

### *sco* cluster retrieval and annotation

*sco* clusters were identified in publicly available genomes of 550 fungi (Table S1) as previously described (Gluck-Thaler & Slot 2018). Briefly, all hits to a previously functionally characterized SCO amino acid query (accession: EAA32528.1) were retrieved from predicted proteomes using BLASTp from the BLAST suite v.2.2.25+ (Altschul *et al.* 1990) with cutoffs (≥30% identity, bitscore ≥50, evalue ≤1e^−5^, and target sequence length 50-150% query length). Additional unannotated sequences were retrieved by tBLASTn of genome assemblies. All genes located within 20 genes upstream and downstream of all *sco* hits (“neighborhood”) were assigned to homolog groups using OrthoMCL v2 (Fischer *et al.* 2011)with an inflation value of 1.5, and homolog groups with limited phylogenetic distributions were discarded. Clusters were then detected by comparing the phylogenetic distributions of homolog group combinations within neighborhoods to distributions expected under null models of microsynteny evolution. A matrix detailing the presence or absence of homolog groups in each cluster was used to calculate Raup-Crick dissimilarity indices for all pairwise comparisons of clusters using the vegan v.2.4-4 package in R (Oksanen *et al.* 2017). The matrix of dissimilarity indices was then subjected to complete linkage clustering, and any clusters separated by less than the empirically determined distance cutoff of 0.05 units were assigned to the same cluster model. As implemented, Raup-Crick dissimilarity values correspond to the probability that the content of two given clusters is non-identical.

Summary descriptions of each cluster model consisted of those homolog groups found in ≥75% of clusters assigned to that model. A representative sequence from each summary homolog group was used to query genome assemblies whose retrieved clusters lacked that homolog group using tBLASTn. Hits occurring within 6 genes upstream or downstream of an *sco* sequence or a sequence part of an *sco* cluster and whose lengths were between 50-150% of the query were manually assembled and considered clustered with that *sco*. All detected gene clusters were re-classified as belonging to a given model if they were missing at most 1 homolog group from that model’s summary. Clusters not meeting this criteria but with at least 1 homolog group from the model’s summary were designated as ‘partial’; clusters meeting the criteria for multiple models were designated as ‘hybrid’; clusters meeting the criteria for full membership in one model and partial membership in another were designated as ‘partial-hybrid’. *sco* homologs not part of any cluster were designated ‘singletons’.

Annotations of all clustered proteins were computed using eggnog-mapper (Huerta-Cepas *et al.* 2017) based on fungal-specific fuNOG orthology data (Huerta-Cepas *et al.* 2016b), and consensus annotations for homolog groups were derived by selecting the most frequent annotation among all members of the group. Conservation of synteny between select clustered regions was assessed with pairwise tBLASTx searches and visualized with EasyFig v2.1, with minimum hit length of 100bp and maximum e-value of 1E-4 (Sullivan *et al.* 2011).

### Phylogenetic analysis and visualization

Preliminary phylogenetic analyses of key clustered genes (*sco*, *gdo, vao* and *dhbd*) and conserved housekeeping genes (*rpb1, rpb2, pol1, hsp60* and *ef1a*) were conducted as follows: all hits to various amino acid queries were retrieved using BLASTp, where hits had a minimum 1e^−5^ evalue cutoff (Altschul *et al.* 1990). Genomes without a hit to a given query were manually searched with tBLASTn, and resulting hits compiled with the rest. All sequences were aligned with mafft v7.221 using the --auto strategy(Katoh & Standley 2013) and the resulting sequence alignment was trimmed with TrimAl v1.4 using the -automated1 strategy (Capella-Gutierrez *et al.* 2009). A preliminary phylogenetic analysis was then performed with FASTTREE v.2.1.10 (Price *et al.* 2010), where trees were mid-point rooted prior to subsequent pruning.

For select clustered genes, a representative sequence was chosen from each major clade in the preliminary midpoint-rooted FASTTREE and used to retrieve the top 10 bacterial hits from the NCBI nr database (last accessed 03/01/2018). These bacterial sequences were combined with the original set of fungal sequences and submitted for preliminary phylogenetic analysis as above. The resulting trees were rooted based on the placement of bacterial sequences. All sequences descending from the last common ancestor of sequences in *sco* clusters were then extracted, and submitted to preliminary phylogenetic analysis as above. The best fitting model of protein evolution for phylogenetic reconstruction was determined according to the AICC using prottest v3.4 (Darriba *et al.* 2011). Trimmed alignments were submitted to RAxML v8.2.9 for maximum likelihood (ML) phylogenetic analysis where 100 rapid bootstraps were used to map support values on the best-scoring ML tree (Stamatakis 2014). For housekeeping genes, the clade of single copy sequences containing the query on the preliminary FASTTREE was extracted and submitted to ML analysis, as above. The majority rule consensus tree of the five housekeeping gene trees was computed in RAxML, and its topology was used to constrain another ML analysis of the trimmed *rpb2* alignment in RAxML with 100 rapid bootstraps.

To quantify the phylogenetic and taxonomic diversity of each key clustered gene phylogeny, we calculated the percentage of total branch length on the *rpb2* species tree for each internal node, as well as the number of taxonomic classes associated with the set of genomes descending from each node.

The trimmed *sco* alignments were additionally used for Bayesian inference with MrBayes v3.2.6 (Ronquist *et al.* 2012). Two independent Metropolis coupled Markov Chain Monte Carlo analyses (options: 4 chains, temp=0.03, starttree=random, prset aamodelpr=mixed, lset rates=invgamma Ngammacat=4) ran until the average standard deviation of split frequencies fell below 0.05, which occurred after 10 million generations. Trees were sampled every 100 generations, and the first 35% of trees were discarded as burn-in. The 50% majority rule consensus tree was obtained using the sumt function.

All phylogenetic trees were visualized using ETE v3(Huerta-Cepas *et al.* 2016a). Tanglegrams were drawn using the ‘ape’ package’s ‘cophyloplot’ function in R (Paradis *et al.* 2004).

### Evolutionary transition analyses

All ancestral state reconstruction analyses were conducted by submitting the 50% credible set of *sco* trees from the MrBayes analysis (minus burnin; see above) and a trait matrix indicating each *sco* sequence’s clustered state (either model 1, model 2, model 3, or unclustered; hybrid clusters were assigned to multiple states) to a BayesTraits v3 multistate MCMC analysis (options: reverse jump with a gamma prior for the rate coefficients whose mean and variance were both seeded from uniform hyper-priors ranging from 0-20; 1.2 billion total iterations; 20% discarded as burnin; sample every 192,000 iterations) (Pagel & Meade 2006). Because our model consists of 12 parameters corresponding to transition rates between each possible clustered state, we used reverse jump MCMC in order to decrease the number of rate parameters to be estimated and decrease the complexity of our model. Reverse jump MCMC integrates all results over model space though an MCMC search, weighing models with different numbers of estimated parameters by their probability, resulting in a posterior distribution of simplified models that best explain the data. Values for the prior and hyper-prior were chosen by examining trace plots of the posterior distributions of various preliminary analyses run with multiple alternative values, and selecting the values that resulted in distributions that did not appear to be prematurely cutoff. Variation in the quality of genome assemblies was accounted for by treating unclustered *sco* singletons located within 6 gene models of a contig or scaffold end as belonging to all clustered states with equal probability. Transition rates were estimated over the entire tree. Transitions between clustered states and gains in clustered state were inferred when the median probability of the ancestral clustered state at a given node increased by ≥75% compared to the median probability of that state at the most recent parental node with ≥0.95 posterior probability support. Transition and gain events that may have occurred over multiple nodes but are supported at a particular node by extant cluster distributions are referred to as having “ambiguous” placement. Due to ambiguity in the ancestral clustered state at/near the root of the tree, we refrained from inferring gains or transitions in clustered states at up to 3 nodes away from the root.

### Testing hypotheses of cluster gains and transitions

In order to obtain an appropriate ML *sco* phylogeny for hypothesis testing, we submitted the trimmed alignment of *sco* sequences for ML analysis, as above, with the added constraint that the topology exactly match that of *sco*’s Bayesian consensus tree (Table S2). It was necessary that the topology of the ML tree to be used for testing alternate hypotheses match that of the Bayesian consensus because the latter was used to reconstruct ancestral states. In order to then assess support for the independence of cluster transitions and origins within a ML framework, we retrieved the per-site log likelihoods of the Bayesian topology-constrained and hypothesis-constrained ML *sco* phylogenies with RAxML (option: -f G) and submitted them to CONSEL v0.2 for analysis with the approximately unbiased (AU) test (Shimodaira & Hasegawa 2001; Shimodaira 2002). The AU test calculates the probability that a given topology has the largest observed likelihood among the set of supplied phylogenies. The null hypothesis that a given phylogeny had the largest observed likelihood was weakly rejected at α ≤ 0.10 and strongly rejected at α ≤ 0.01. To obtain constrained ML trees, the trimmed alignment of *sco* sequences was submitted to RAxML (options: -f d, -g) along with a constraint tree specifying the monophyly of particular sets of sequences (see Table S3 for all constraint criteria). These trees were non-comprehensive (i.e., they only contained sequences explicitly mentioned in the hypothesis), and besides the bifurcating node specifying the monophyletic constraint, consisted only of multifurcating nodes. The placement of taxa not present in the constraint tree and multifurcations were resolved using a ML search. Each constrained ML phylogeny was manually verified to adhere to the specified constraint.

### Molecular Cloning and Protein expression

An oligo-dT-primed cDNA library constructed using SuperScript^®^ III First-Strand Synthesis System for RT-PCR (#18080-051, Invitrogen Life Technologies, Waltham, MA, USA) was used as the template for obtaining the coding sequences of four fungal SCOs (*Dssco* from *Diplodia sapinea* (MH350427); *Mosco* from *Magnaporthe oryzae* (MH350428); *Pasco* from *Podospora anserina* (MH350429); *Prsco* from *Penicillium roqueforti* (MH350430)) by PCR amplification using Platinum Taq DNA polymerase (#11304-011, Invitrogen, Waltham, MA, USA). Specially designed oligonucleotide primer pairs for enzyme-free cloning into pETite C-His Kan Vector (#49002- 1, Expresso T7 Cloning Kit, Lucigen Corp., Middleton, WI, USA) for C-terminal His tag protein expression and cell lysate preparations were used. The oligonucleotide primer pairs used in the present study are listed in Table S4. The resulting recombinant plasmids were transformed into the HI-Control 10G host strain and following sequence verification freshly transformed into HI-Control BL21 (DE3) chemically competent cells for expression and cell lysate preparation, respectively, following manufacturer’s instructions. Sequence verified transformant(s) were grown in 5 mL of Luria-Bertani (LB) liquid medium containing 50 µg/mL kanamycin for 16 h at 37 °C with 225 rpm shaking. Cultures were transferred to 100 mL of fresh LB liquid media in a 500 mL flask and incubated at 37°C until cultures reached an optical density (OD) at 600 nm of 0.4–0.6. Protein expression was induced with 1 mM isopropyl β-D-1-thiogalactopyranoside (IPTG, #I3301, Teknova, Hollister, CA, USA) for 4 h at 22–25°C. Later, cells were pelleted, and lysed using 1 mg/mL lysozyme (#3L2510, Teknova, Hollister, CA, USA) and incubated on ice for 30 min. Soluble proteins were collected after centrifugation of cell lysates at 10,000 x g for 30 min at 4 °C.

### *In vitro* enzymatic assays

All reagents and analytical grade substrates (resveratrol, pterostilbene, piceatannol, piceid, isorhapontigenin, isoeugenol, 3,5-dihydroxybenzaldehyde, 3,5-dimethoxybenzoic acid, 3,4-dihydroxybenzaldehyde and 4-hydroxybenzaldehyde) used for enzymatic assays were purchased from Sigma-Aldrich (St. Louis, MO, USA), while pinosylvin (CAS # 22139-77-1) was purchased from Sequoia research products (Pangbourne, UK). Briefly, assays with respective cell lysates (≤ 20 µg) were carried out in 100 µL incubation buffer containing 1mM Tris(2- carboxyethyl) phosphine hydrochloride (TCEP, #C4706), 0.3mM FeSO4 (#215422), 100 mM HEPES (#H3375) and 1 mg/mL Catalase (#C1345). Control incubations were performed by preparing cell lysates of empty HI-Control BL21 (DE3) chemically competent cells. The cell lysate (*c*. 15-20 µL) was mixed with the incubation buffer and allowed to stand for 5 min to equilibrate. Substrates were dissolved in DMSO (#D128, Fisher Scientific, Columbus, OH, USA) and added to a final concentration of 100 µM. Reactions were incubated at 27°C for 30 min in the dark and extracted with 2x 100 µl of ethyl acetate (#E195, Fisher Scientific, Columbus, OH, USA) containing 1.0 mg/ml butylated hydroxyanisole as an internal standard (BHA, # B1253). Finally, the supernatants were combined and dried in an Eppendorf vacufuge concentrator (#07-748-13, Fisher Scientific, Waltham, MA, USA) for 25 min. All samples were re-suspended in 100 µl of pure HPLC grade methanol (#A456, Fisher Scientific, Columbus, OH, USA), vortexed, and placed in a water bath sonicator for 10 minutes before being analyzed by UPLC.

### UPLC separation and analyses

Products of the in vitro enzymatic reactions were analyzed by UPLC, and relatively quantified by using BHA (Figure S1). For UPLC measurements, the column used was Cortecs UPLC C18 (2.1 < 100 mm. 1.6 µm, #186007095, Waters Corp., Milford, MA, USA) with a column temperature of 40 °C and sample tray held at 10 °C. The mobile phase consisted of 0.1% acetic acid (A, aqueous) and acetonitrile (B) with a flow rate of 0.3 mL/min throughout. The gradient used was as follows: 0–0.25 min of 10 % B, 0.25–2 min of 10–27 % B, 2–4 min of 27-50 % B, 4–8 min of 50–90 % B, 8.01–10 min of 90 % B, and 10.01–12 min of 10 % B again. Area (µV*sec) for each peak was recorded and cleavage product concentration was calculated using external standard equation (C_x_) = A_x_ / CF_ave_, where A_x_ is the area of the analyte and CF_ave_ is the calibration factor. CF_ave_ was calculated using the formula CF = Area of the analyte / Conc. of the internal standard. Percent substrate consumption and relative enzymatic activity were calculated based on area (µV*sec) of the substrate peaks in the chromatogram before (100µM) and after enzymatic reactions.

### Motif analysis of protein sequences

Previously identified substrate binding motifs(McAndrew *et al.* 2016) were retrieved by aligning two previously characterized bacterial SCOs with all 9 characterized fungal SCOs (5 described in previous studies, 4 described here), and analyzing regions of interest using commandline MEME v.4.12.0 (Bailey & Elkan 1994). Identified motifs were retrieved from the complete set of scos using FIMO in the MEME suite.

## Results

### *sco* homologs are alternately located in three distinct gene clusters

Using a previously described cluster detection method (Gluck-Thaler & Slot 2018), we identified unexpectedly retained synteny in the gene neighborhoods surrounding 466 monophyletic *sco* homologs retrieved from 550 fungi, and found that *sco* variably forms clusters with three distinct groups of genes, which we refer to as gene cluster models (Methods; Figure 1, Figures S2-5, Table S5). Model 1 contains four genes: *sco*, a 2,3-dihydroxybenzoate decarboxylase (*dhbd;* EC# 4.1.1.46), an aldehyde dehydrogenase (EC# 1.2.1.10), and a gene with unknown function containing a Snoal-like domain (PF13577) and part of the NFT2-like superfamily (SSF54427). Model 2, which was previously identified by Greene et al. (2014), contains five genes: *sco*, a gentisate dioxygenase (*gdo;* EC# 1.13.11.4), a salicylate hydroxylase (EC# 1.14.13.1), a fumarylpyruvate hydrolase (EC# 3.7.2.5), and a Zn(2)-Cys(6) transcription factor. Model 3 consists of two genes: *sco*, and a vanillyl alcohol oxidase (*vao;* EC# 1.1.3.38). A total of 80 model 1 clusters, 35 model 2 clusters, 55 model 3 clusters and 1 hybrid model 2+3 cluster were found distributed among 120 Pezizomycotina genomes. Of these, 4 model 1 clusters, 1 model 2 cluster, and 29 model 3 clusters contained at least one gene from other cluster models (but not enough to be considered true hybrids). 31 genomes harbored more than one type of cluster model. No genome was found to contain more than one copy of either model 1 or model 2 clusters; however, 8 genomes contain multiple model 3 clusters. A total of 52, 28, and 0 *sco* homologs were part of partial model 1, model 2 and model 3 clusters, respectively (defined as having <75% of the genes in the cluster model), while 203 *sco* homologs were not identified as clustered according to our criteria (see Methods). We found two additional model 2 clusters in the carT clade of fungal carotenoid cleavage dioxygenases when we expanded our search for cluster models outside of the monophyletic *sco* clade (Figure S2).

**Figure 1.**
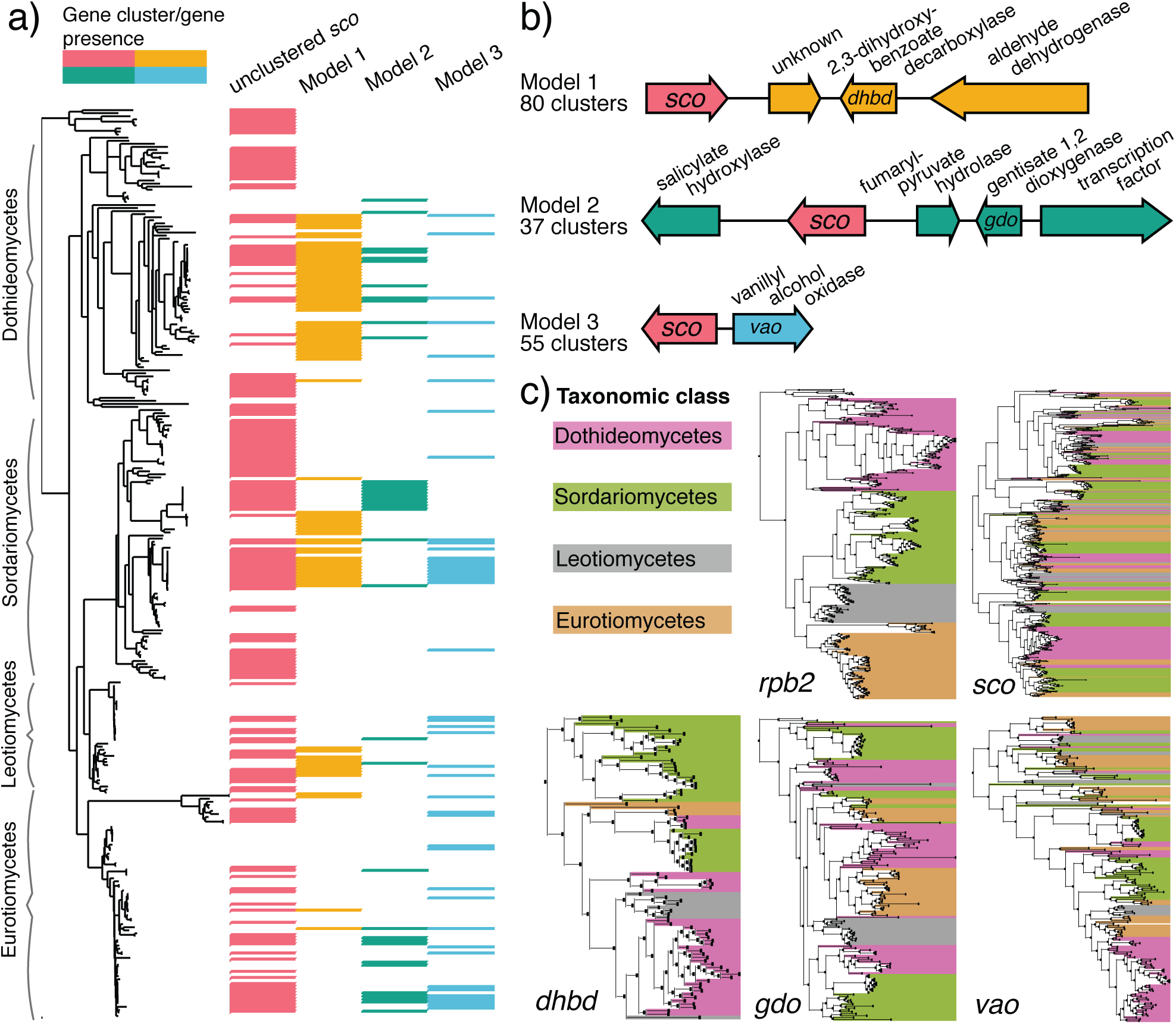
The distributions of three types of stilbene cleavage oxygenase (*sco)* gene clusters in fungi. a) A rooted, constrained maximum likelihood phylogeny based on the amino acid sequences of the second largest subunit of RNA polymerase II (*rpb2*; Methods) representing relationships among all 288 Pezizomycotina genomes examined in this study and 9 outgroup fungal genomes. Four major Pezizomycotina classes are indicated to the left. This phylogeny contains all 212 genomes that have at least 1 *sco* homolog (of which 203 are from the Pezizomycotina) from a database of 550 genomes. Matrix indicates presence/absence of unclustered *sco* genes and three cluster models in individual genomes, color-coded by gene/cluster model type. b) Representative example of each *sco* cluster model with genes color-coded according to model type (except *sco*, which is colored red): model 1: *Ophioceras dolichostomum* (Sordariomycetes), Ophdo1_1340.90_c1; model 2: *Magnaporthe oryzae* (Sordariomycetes), Maggr1_117659_c2; model 3: *Podospora anserina* (Sordariomycetes), Podan2_2307_c1 (Table S5). Predicted functions are indicated, and key genes in each model are labeled (*dhbd*=2,3-dihydroxybenzoate decarboxylase; *gdo*=gentisate 1,2 dioxygenase, *vao=*vanillyl alcohol oxidase). c) Rooted maximum likelihood phylogenies of *rpb2*, *sco*, *dhbd*, *gdo*, and *vao* amino acid sequences with associated taxa color-coded by Pezizomycotina class. All color codes are consistent within and across figures.

Each cluster model is defined by a single key enzyme-encoding gene with homologs known to function in monophenolic catabolism. *dhdb* (model 1) is in the KEGG orthology group K14333, which also contains a characterized enzyme from *Aspergillus oryzae* (accession: P80346.1) that catalyzes the decarboxylation of 2,3-dihydroxybenzoate to catechol. Fungal *dhbd* sequences additionally share a conserved amidohydrolase domain (PF04909) with *Sphingobium* sp. SYK-6 *ligW2* (accession: WP_014075111.1), an enzyme that participates in the degradation of lignin-derived biphenyl compounds through the non-oxidative decarboxylation of 5-carboxyvanillate (Peng *et al.* 2005). *gdo* (model 2) is in the KEGG orthology group K00450, contains a conserved cupin-fold (PF07883.10), and is related to bacterial *gdo*s that participate in gentisate degradation by catalyzing the ring cleavage of gentisate to 3-maleylpyruvate (Werwath *et al.* 1998). *vao* (model 3) contains FAD-binding (PF01565.22) and FAD-linked oxidase (PF02913.18) domains, and includes the only functionally characterized fungal *vao* from *Penicillium simlissisum, vaoX* (accession: P56216). *vaoX* catalyzes the conversion of vanillyl alcohol to vanillin, but is also active on 4-hydroxybenzyl alcohols, 4-hydroxybenzylamines, 4-(methoxy-methyl)phenols, and 4-allylphenols (Fraaije *et al.* 1995).

### SCOs encoded in different clusters have conserved functions and substrate binding motifs

Fungal SCOs share a number of secondary structural and functional residue features with the characterized bacterial NOV1 (McAndrew *et al.* 2016). Fungal SCOs contain the conserved protein domain RPE65 (pfam03055), which defines members of the carotenoid cleavage dioxygenase family in plants and bacteria, and SCOs in bacteria and fungi. We identified previously characterized residues associated with tetradentate iron coordination in the SCO active site and with the essentiality of 4’-OH groups for stilbene cleavage by bacterial SCO (Figure 2, Table S6) (Harrison & Bugg 2014; McAndrew *et al.* 2016). Four histidine residues and three glutamate residues in SCO contribute to the 4-His + 3-Glu (second shell glutamate) dual-sphere metal binding catalytic center involved in canonical non-heme iron (II) dependent CCD-type enzymes (Harrison & Bugg 2014). These residues are conserved in all nine functionally characterized SCO sequences, and amino acid motifs indicate additional imperfect conservation in surrounding residues. 60% of all retrieved fungal SCOs were conserved in all these residues (Table S7). The motifs in the conserved 4-His + 3-Glu dual-sphere metal binding catalytic center are: (F/M)(T/C)AHPK*X*, *X*(M/F)(I/M)HD(C/F)(A/G), (F/M)*XX*H(T/V)*XX*, *X*(G/Q)*X*HG*X*(W/F), A*X*KE*XXX*, *XXX*EF*X*(R/Q), and *XX*(Q/G)E(P/C)*XX*, respectively.

**Figure 2.**
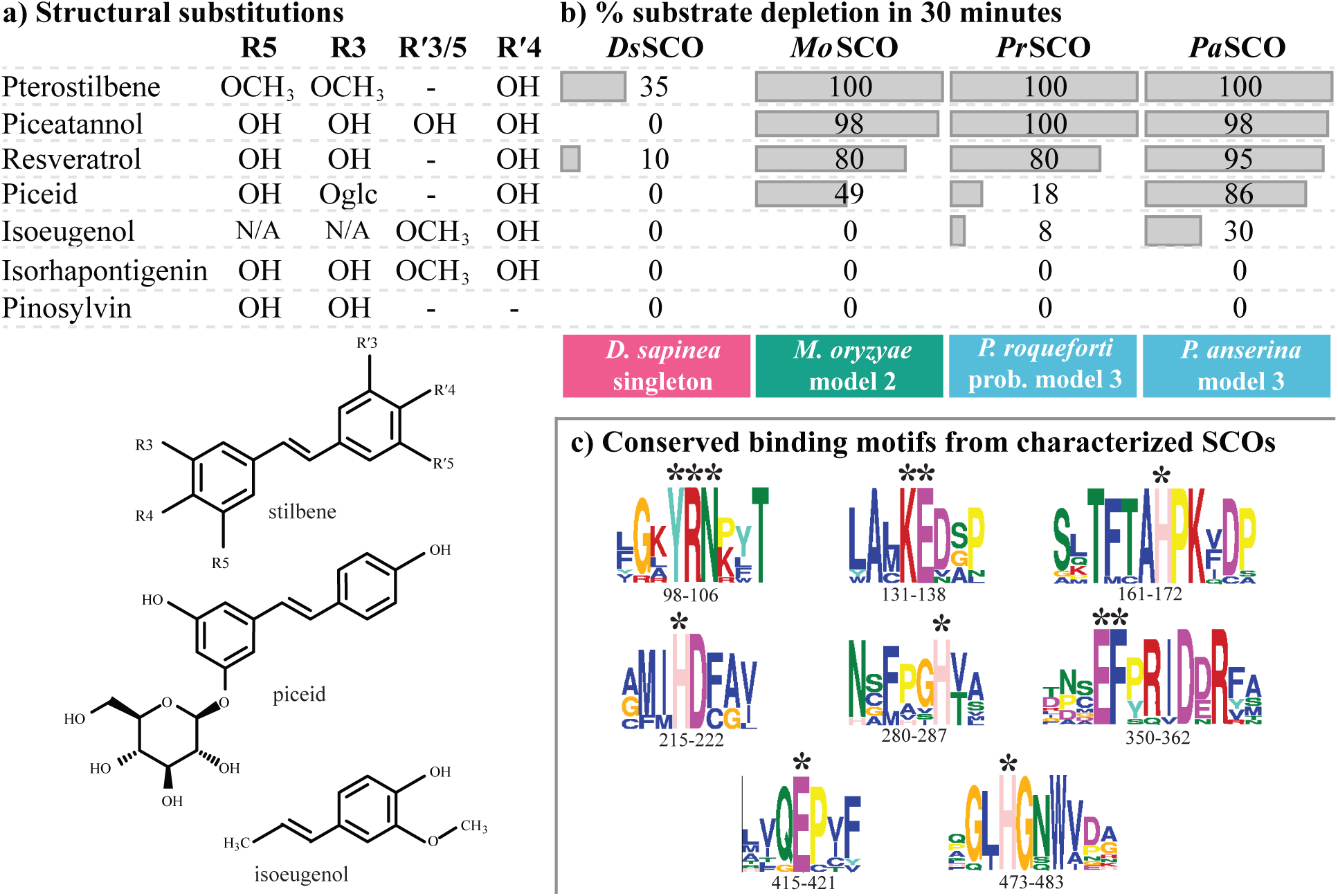
Biochemical characterization of four fungal stilbene cleavage oxygenase (SCO) enzymes encoded in alternate cluster types. a) Chemical structures of all seven molecules used for enzyme characterization analyses. b) Percent depletion of the seven examined molecules after a 30-minute incubation with each of four heterologously expressed fungal SCO enzymes encoded in different cluster models: *Ds*SCO from *Diplodia sapinea* (accession: MH350427; not part of any identified cluster models); *Mo*SCO from *Magnaporthe oryzae* Guy11 (accession: MH350428; model 2); *Pr*SCO from *Penicillium roqueforti* FM164 (accession: MH350430; probable model 3); *Pa*SCO from *Podospora anserina* S mat+ (accession: MH350429; model 3). c) Eight conserved substrate-binding motifs from two characterized bacterial SCO sequences and all nine characterized fungal SCO sequences (four from this study, five from previous studies, Table S6 and S9). Residues of interest (indicated positions relative to NOV1 from *Novosphingobium aromaticivorans*, accession YP_496081.1) discussed in the main text are indicated with an asterisk (*), and include the YRN and KE motifs involved in the recognition and deprotonation of 4’-OH groups on stilbene molecules, the EF motif involved in hydrogen bonding with the 3/5 hydroxyl group of resveratrol molecules, and the 4-Histidine(H) + 3-Glutamic acid(E) dual-sphere metal binding catalytic center found in non-heme iron (II) dependent carotenoid cleavage dioxygenases.

Similarly, Tyr and Lys residues essential for recognition and deprotonation of 4’-OH group of stilbenes in NOV1, are fully conserved in the nine functionally characterized proteins (Figure 2c). The Tyr residue is part of a YRN motif fully conserved in the characterized protein, and in approximately 86% of all retrieved fungal SCOs, and the Lys residue is part of a KE motif fully conserved in characterized proteins, and in approximately 82% of all retrieved fungal SCOs (Table S7). Finally, a Glu residue previously shown in NOV1 to hydrogen bond with the 3/5 hydroxyl groups of resveratrol (McAndrew *et al.* 2016), is fully conserved in an EF motif (Figure 2c) in the characterized proteins, and present in approximately 82% of all fungal homologs (Table S7). Finally, the Ser residue (S283) in NOV1, speculated to be involved in dioxygenase activity on isoeugenol, a non-stilbene possessing a 4’-OH group, was not found in fungal SCOs that cleave isoeugenol (Table S6).

Four SCOs encoded in different genomic contexts in four different fungal species were profiled for substrate preferences (Figure 2, Figure S5, Table S8): (1) *Ds*SCO (unclustered) from the pine necrotroph *Diplodia sapinea*; (2) *Mo*SCO (model 2 cluster) from the grass hemibiotroph *Magnaporthe oryzae*; (3) *Pa*SCO (model 3 cluster), from the dung saprotroph *Podospora anserina*; (4) *Pr*SCO (probable model 3 cluster, see below) from the cheese-associated saprotroph *Penicillium roqueforti*. Although *Prsco* is provisionally classified as unclustered, we suspect this may be due to the fragmented nature of its genome, and that it may actually be located in a model 3 cluster with a *vao* sequence. Both *sco* and *vao* in *P. roqueforti* are terminal genes on their respective contigs, and are located within co-diversifying monophyletic clades of sequences found in model 3 clusters (Figure S6). Additionally, close relatives of *P. roqueforti*’s *sco* and *vao* in *Penicillium glabrum* and *Penicillium chrysogenum* are found together in model 3 clusters. No SCOs from model 1 were included, as the above 4 SCOs were selected and characterized before the development of our most recent cluster detection algorithm that led to the discovery of the model 1 cluster type. However, the previously characterized SCO from *C. globosum* is encoded in a partial model 1 cluster (Figure S5). All *sco*s were expressed under standard growth conditions, except for *Dssco,* which was only expressed when induced with 0.2 % resveratrol or ground Austrian pine needle agar medium (Luchi *et al.* 2007).

Substrate cleavage specificities of heterologously expressed proteins were similar among the four enzymes in a panel consisting of the stilbenes resveratrol, pterostilbene, piceatannol, piceid, pinosylvin, and isorhapontigenin, and the monophenolic isoeugenol, with some notable exceptions (Figure 2 and Table S8). Pterostilbene is the preferred substrate of all four enzymes, generally followed by piceatannol, and then resveratrol. None of these enzymes was active on isorhapontigenin, which differs from piceatannol by methylation of the R’3/5-OH, or pinosylvin, which lacks a 4’-OH group. All of these results are consistent with cleavage of more limited panels of stilbenes by the other 5 characterized fungal SCOs (Brefort *et al.* 2011; Díaz-Sánchez *et al.* 2013), and similar to that of the extensive panel of stilbenes and related molecules tested on bacterial NOV1 (McAndrew *et al.* 2016)(Table S9).

We identified a number of differences among these SCO enzymes in terms of substrate diversity and level of activity under the experimental conditions (Figure 2). Of all the four tested enzymes, *Pa*SCO had the highest activity and broadest range in cleavage profiles, effectively cleaving five out of six stilbenoid compounds and isoeugenol. *Pr*SCO had the same range of substrates as *Pa*SCO, but was not as effective at cleaving the non-stilbene isoeugenol or piceid, a glycosylated derivative of resveratrol whose cleavage has not been previously reported. *Mo*SCO had similar affinities for stilbenoid compounds to PrSCO but did not cleave isoeugenol. *P. anserina*, *P. roqueforti,* and *M. oryzae* SCOs all had high cleavage activity on pterostilbene, piceatannol and resveratrol, but compared to the other enzymes, *Pa*SCO was much more active on piceid (Table S8). *Ds*SCO was somewhat active on pterostilbene and resveratrol, but no other compounds tested. Interestingly, in the very closely related but less virulent pine pathogen *D. scrobiculata*, *sco* is a pseudogene. The greatest differences in activity among the different enzymes were observed in isoeugenol and piceid.

### The evolution of *sco* cluster genes suggests complex evolutionary histories, and co-diversification within clusters

In a maximum likelihood phylogeny of bacterial, plant and fungal carotenoid cleavage dioxygenases and SCOs, all fungal SCOs are monophyletic with previously characterized bacterial SCOs, to the exclusion of bacterial, plant and fungal carotenoid cleavage dioxygenases (EC# 1.13.11.59) involved in carotenoid metabolism (Figure S2). Carotenoid cleavage dioxygenases fall into two distinct clades (carX and carT) each containing groups of fungal sequences that are monophyletic with earlier diverging bacterial and plant sequences. Maximum likelihood phylogenies of *sco* and key genes from each of the *sco* cluster models (*dhbd* from model 1: Figures S7-8; *gdo* from model 2: Figures S9-10; *vao* from model 3: Figure S10-12) in fungi portrayed complex patterns of inheritance compared to the conserved housekeeping gene *rpb2*, which closely tracks accepted species relationships (Figure 1c, Figure S13). In the *sco* and *vao* gene phylogenies, sequences belonging to fungi from different taxonomic classes are frequently found as sister taxa, while the phylogenies of *gdo* and *dhbd* appear to conflict somewhat less with vertical inheritance among species. On average, taxa descending from each node on the maximum likelihood *sco, vao, gdo* and *dhbd* phylogenies cover 0.051%, 0.054%, 0.047% and 0.035% of the *rpb2* tree and are distributed across 1.56, 1.61, 1.30 and 1.19 taxonomic classes, respectively (Table S10). For comparison, taxa descending from each node on the *rpb2* phylogeny cover 0.03% of the *rpb2* tree and represent 1.07 taxonomic classes, on average.

We compared clades of key genes from each of the three cluster models with the *sco* topology to investigate the patterns of clustered gene co-diversification (Figure 3). Large clades of clustered *dhbd* (model 1) and *gdo* (model 2) sequences co-diversify with their neighboring *sco* sequences, suggesting that clustered sequences in the respective gene families have been co-inherited since they became clustered early on. Although clades of clustered *vao* sequences also co-diversify with their co-clustered *sco* sequences, distantly related *sco* sequences tend to cluster with closely related vao sequences, suggesting multiple and more recent independent origins of their clustering. Congruence between clustered clades on different trees is independent of whether the genes’ topologies are consistent with species trees. Clustered homologs of *gdo* and *vao* are often closely related to unclustered homologs.

**Figure 3.**
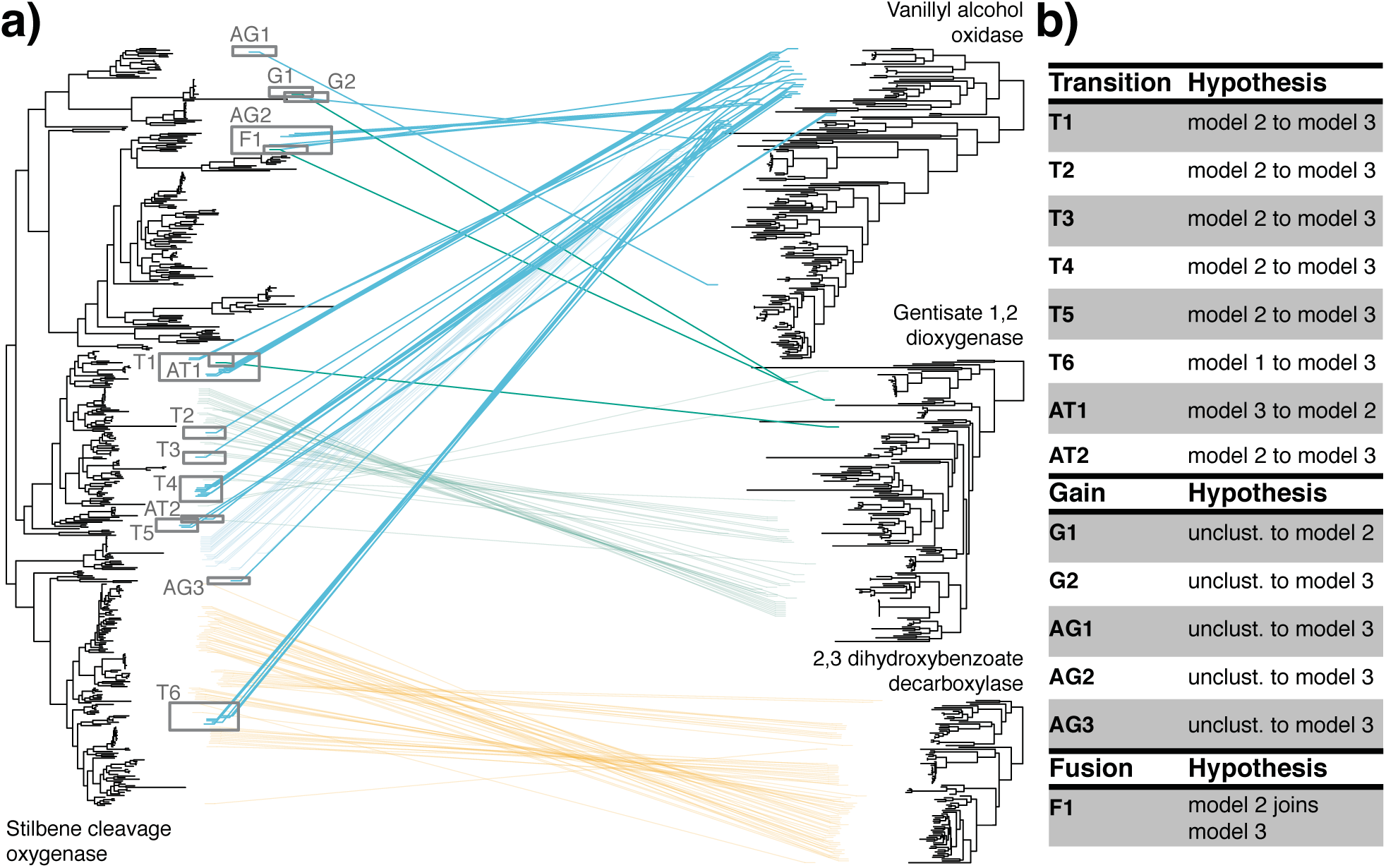
Transitions between and gains of distinct cluster types across the stilbene cleavage oxygenase (*sco*) phylogeny. a) A tanglegram of the 50% majority rule consensus Bayesian *sco* phylogeny and maximum likelihood phylogenies of 2,3-dihydroxybenzoate decarboxylase (*dhbd*; key gene model 1), gentisate 1,2 dioxygenase (*gdo*; key gene model 2) and vanillyl alcohol oxidase (*vao*; key gene model 3) amino acid sequences. Connecting lines indicate genes within the same *sco* gene cluster, color-coded by cluster model. Sequences descending from nodes with inferred transition events between, or gain and fusion events of clustered states are outlined with grey squares (labeled by event ID) and have bold connecting lines. b) Descriptions of cluster transition, gain and fusion events inferred through ancestral state reconstruction (details provided in Figure S5 and Table S11). Transition and gain events with ambiguous ancestral states but supported by extant cluster distributions are prefixed with an ‘A’.

### *sco* underwent multiple transitions to and between cluster models

Ancestral clustered states of *sco* (i.e. models 1-3 or unclustered) were estimated within a Bayesian framework, enabling the inference of transitions between and gains of distinct clustered states while integrating over the posterior distribution of *sco* topologies (Figure 3, Figure S5). Transition rates between clustered states sampled during reverse jump MCMC analysis roughly fall into two categories, those with a median rate coefficient of ∼2 (model 1 to unclustered, model 2 to unclustered, model 3 to unclustered, model 2 to 3) and those with a median rate coefficient of ∼0 (unclustered to model 1, unclustered to model 2, unclustered to model 3, model 1 to model 2, model 2 to model 1, model 1 to model 3, model 3 to model 1, model 3 to model 2) (Figure S14).

We identified 6 distinct events (T1-6) and 2 ambiguously placed events (Methods; AT1-2) in which *sco* homologs are inferred to have transitioned between one cluster model and another (Figure 3, Figure 4, Table S11). Transitions from model 2 to model 3 occurred most frequently (T1-T5, AT2), followed by transitions from model 1 to model 3 (T6) and transitions from model 3 to model 2 (AT1). Interestingly, we also inferred the replacement of a *vao* in a model 3 containing taxon with a more divergent *vao* homolog (Figure S6).

**Figure 4.**
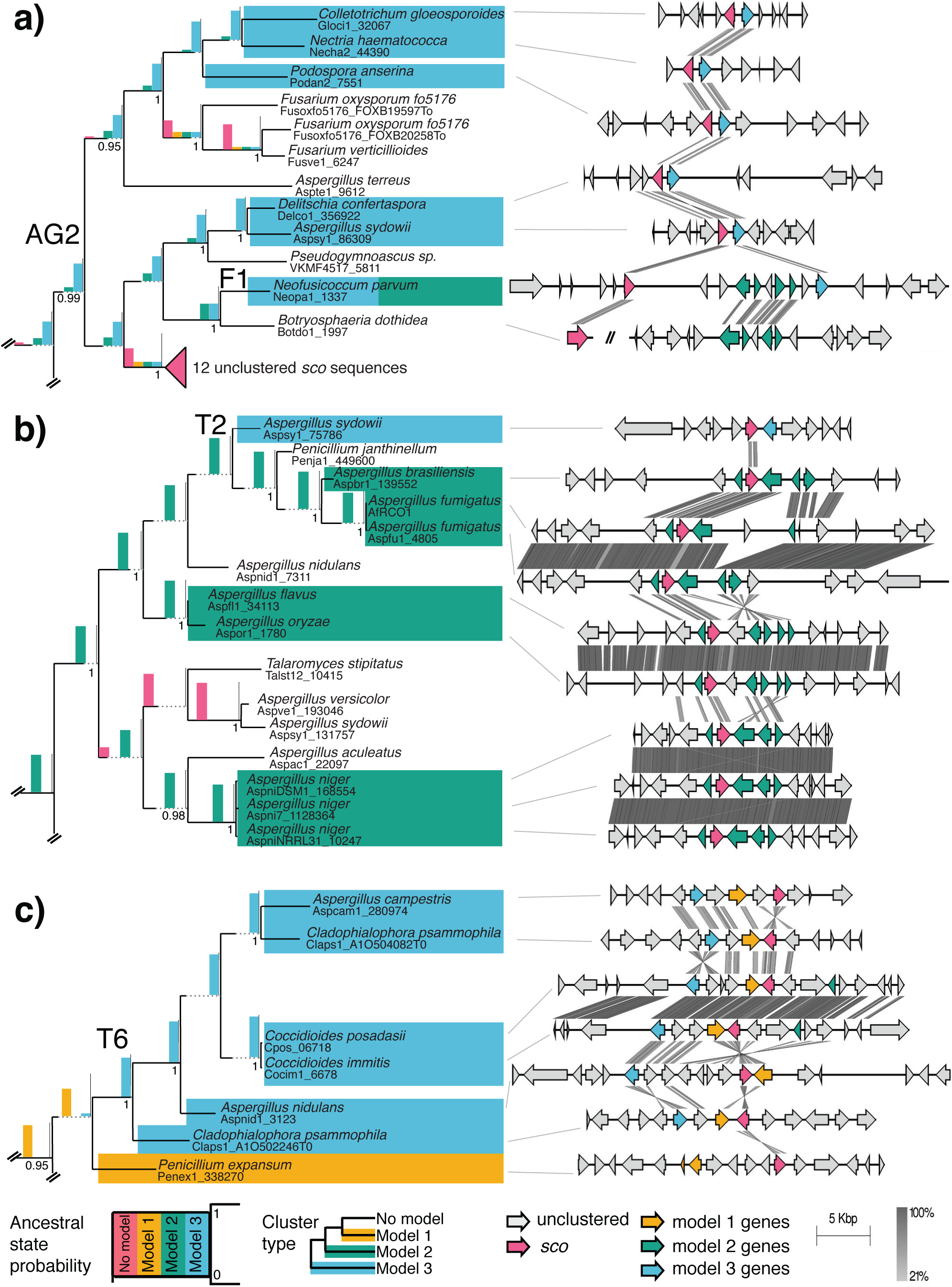
Synteny comparisons within three lineages of stilbene cleavage oxygenase (*sco*) loci inferred to have experienced transition or fusion events between different cluster types. All depicted trees on the left were extracted from the 50% majority rule consensus Bayesian *sco* amino acid phylogeny, and can be viewed in their original context in Figure S5. Tree tips are labeled with binomial species names and protein IDs, and are color coded by the cluster model to which they belong. Transition, gain and fusion event IDs are labeled above the nodes at which they are inferred to have occurred (Figure 3, Table S11). Histograms depicting the median probability of reconstructed ancestral clustered states (ranging from 0 to 1) are drawn above their respective branches. Dotted gray lines artificially extend branches that would otherwise be too short to accommodate histograms. Support values of ≥0.95 posterior probability are drawn beneath their respective branches, while support values <0.95 are not shown. Gene schematics on the right are color coded by the *sco* cluster model to which they belong (if any), except for *sco*, which is always colored red. Shaded-lines drawn between gene schematics indicate percent identity, as determined by pairwise tBLASTx comparisons. Note that genes from different families part of the same cluster model are shaded with the same color. a) Inferred fusion between model 2 and model 3 clusters in *Neofusicoccum parvum* (Dothideomycetes). Note that the close relative *Botryosphaeria dothidea* (Dothideomycetes) has an unclustered *sco* and a model 2 cluster lacking *sco* located on different contigs b) Inferred *sco* transition from model 2 to model 3 cluster type. c) Inferred *sco* transition from model 1 to model 3 cluster type.

We identified one special case of a transition in the form of a cluster fusion (F1) consisting of a single locus containing a *sco* sequence associated with all accessory model 2 and 3 genes in *Neofusicoccum parvum* (Dothideomycetes) (Figure 4). Model 3 is inferred to be the ancestral clustered state of this clade having arisen through a gain in model 3 clustered state (AG2). The *N. parvum vao* homolog (Neopa1_tbn3; detected through tBLASTn) is most closely related to Sordariomycete sequences, while two closely related species, *Macrophomina phaseolina* and *Botryosphaeria dothidea*, contain model 2 clusters lacking *sco* sequences. Coupled with the observation that the *N. parvum sco* homolog (Neopa1_1337) is found in a region of the phylogeny with poor resolution and high taxonomic diversity, these lines of evidence are consistent with the fusion of a vertically inherited model 2 cluster (without *sco*) with ambiguously inherited *sco* and *vao* genes.

We also identified 2 distinct events (G1-2) and 3 ambiguously placed events (AG1-3) where *sco* is inferred to have transitioned from an unclustered state to either a model 3 cluster (G2, AG1-3) or model 2 cluster (G1). Although not included in our ancestral state reconstruction analysis involving the monophyletic *sco* clade, at least one other independent gain of a model 2 clustered state appears to have occurred in the carT clade of carotenoid cleavage dioxygenases involving two sequences (Aspve1_89005 and Lorju1_471378) that are very distantly related to the *sco* clade (Figure S2).

With regards to the SCO enzymes we characterized here, we infer that *Pa*SCO and *Pr*SCO are in a small clade recently descended from T4 (transition from model 2 to model 3), *Mo*SCO is in a clade with an ancient model 2 clustered state, and *Ds*SCO is in a very small clade that recently lost model 3 to become unclustered.

After confirming that the likelihood of the Bayesian topology-constrained *sco* ML tree did not differ significantly from the true optimal ML *sco* phylogeny (p = 0.364, Table S2), we sought to validate inferred transition events within a maximum likelihood framework using constrained topologies corresponding to hypotheses regarding the independence of transition events (Figure 3, Figure S5, Table S3 and S11). We failed to reject the null hypothesis of no transition from model 3 to model 2 among the descendants of node_387_0. 863 (test 4; p = 0. 142), indicating ambiguous support for event AT1 as a true transition event. We failed to reject the null hypothesis that events T3 and T4 have independent origins (test 6; p = 0.195), suggesting that the model 3 clusters descending from these events may have arisen through a single transition event. We also failed to reject the null hypothesis of no independent transitions from model 2 to model 3 at events AT2 and T5 (test 9; p = 0.133), suggesting that model 3 clusters descending from node_547_0.621 may have arisen through a single transition event. We weakly rejected the null hypothesis of no independent transitions among the descendants of node_491_0.663 (test 5; affects events T2 and T3; p = 0.071), of node_859_0.992 (test 12; affects event T6; p = 0.074), and of node_861_0.992 (test 11; affects event AG3, T6; p = 0.079). We strongly rejected the null hypothesis of no independent transitions among the descendants of node_522_0.786 (test 7; affects events T2, T3, T4; p = 0.003), of node_523_1.00 (test 8; affects events T1, T2, T3, T4; p = 3.00E-05), and of node_558_0.968 (test 10; affects events T1, T2, T3, T4, T5, AT2; p = 8.00E-54). We also strongly rejected the null hypothesis of no independent transitions to model 2 clusters from either model 1 or model 3 across the entire *sco* phylogeny (test 2; p = 1.00E-5), and the null hypothesis of no independent transitions to model 3 clusters from either model 2 or 3 across the entire *sco* phylogeny (test 3; p = 8.00E-8). Together, these constraint analyses support a scenario where the *sco* gene family has experienced at least 4 transition events and 1 ambiguous transition event during the course of its evolutionary history.

## Discussion

Fungi are consummate decomposers, and their catabolic activities often underlie their saprotrophic, mutualistic and pathogenic interactions with plants. Many plants produce SMs that inhibit fungal growth, and correspondingly, the ability of fungi to degrade plant SMs like stilbenes can affect how they colonize plant tissues (Michielse *et al.* 2012; Hammerbacher *et al.* 2013; Kettle *et al.* 2015). Studies on the diversification of biosynthetic SM pathways in plants have received considerable attention in recent years (Wisecaver *et al.* 2017; Moghe *et al.* 2017; Leong & Last 2017), yet little is known about the processes underpinning the co-evolution of cognate degradative pathways in fungi. One fungal strategy to degrade plant-produced stilbenes involves the deployment of SCO, a family of fungal enzymes that degrade stilbene compounds (Brefort *et al.* 2011; Díaz-Sánchez *et al.* 2013). Although the pathways in which SCO participates are yet to be described, we report here that *sco* homologs are alternatively organized into three distinct types of MGCs, leading us to ask how different evolutionary processes may have affected the evolution of stilbene catabolism in fungi. In general, metabolic diversification can proceed through combinatorial evolution, where new combinations of existing enzymes coalesce into new pathways, and through sequence-based evolution, where changes to gene coding sequences result in new enzymatic activities (Wagner 2011). Our biochemical and phylogenetic investigation suggests that combinatorial evolution plays a primary role in promoting the diversification of stilbene degradation pathways in fungi, while *sco*’s enzymatic function is more constrained.

### Convergent assembly of *sco* clusters suggests selection for specific gene combinations

The generation of new phenotypes through combining existing genotypes is a key driver of metabolic diversity (Takiguchi *et al.* 1989; Wagner 2011). MGCs participating in the biosynthesis and degradation of SMs are increasingly recognized as hotspots for combinatorial evolution (Copley 2009; Lind *et al.* 2017; Gluck-Thaler & Slot 2018). These loci experience frequent gene pseudogenization, duplication, insertion, and deletion events that differentiate gene content even amongst closely related species (Reynolds *et al.* 2017; Lind *et al.* 2017). Several MGCs are inferred to have arisen through the relocation of genes within the same genome (Proctor *et al.* 2009), while others have arisen through relocation and fusion with existing MGCs (Lind *et al.* 2017), similar to the cluster fusion we observed here in *N. parvum* (Figure 4). Many MGCs are located in recombination hotspots, such as subtelomeres (Wu *et al.* 2009; Croll *et al.* 2015) and TE-rich regions (Fleetwood *et al.* 2007; Lind *et al.* 2017), which may help generate combinatorial diversity at these loci.

In this study, we provide evidence that combinatorial evolution has impacted the evolutionary trajectory of stilbene catabolism. We observed multiple independent instances in which *sco* genes shifted genomic context either from an unclustered state to a distinct cluster type or between cluster types, suggesting convergent selection for specific gene combinations (Figure 3, Figure 4, Table S11). Convergence is indicative of strong directional selection for optimal phenotypes in similar environments (Conant & Wagner 2003; Bittleston *et al.* 2016). Several other examples of convergent evolution of particular combinations of clustered genes have previously been reported, notably galactose utilization clusters in divergent yeast lineages (Slot & Rokas 2010), cyanate detoxification clusters in the Pezizomycotina (Elmore *et al.* 2015), nicotinate degradation clusters in fungi and bacteria (Amon *et al.* 2017), and triterpene biosynthesis clusters in plants (Field & Osbourn 2008). While our analysis was limited to examining cluster assembly from the perspective of *sco*, the organizational histories of genes associated with *sco* are also marked by convergence. For example, the *sah*, *fph,* and *gdo* genes present in model 2 clusters have convergently clustered without *sco* in bacteria and fungi (Liu *et al.* 2011).

One outstanding question regarding convergent cluster assembly is why clustering is selected at all. Although selection for coordinated expression is often proposed as a driving force behind clustering, support for this hypothesis is mixed: an explicit test using galactose utilization clusters in yeast suggested that co-expression is a consequence but not necessarily a cause of clustering (Lang & Botstein 2011), while homologs of clustered fungal metabolic genes are more likely to be co-expressed in primates suggesting a causal relationship (Eidem *et al.* 2015). Whether or not coordinated expression is selectable may ultimately depend on the fitness consequence of sub-optimal regulation, which is likely more severe if pathway intermediates are toxic. Clustered gene pairs are more likely to encode enzymes that are linked by a toxic intermediate (McGary *et al.* 2013), and some of the predicted intermediary metabolites of stilbene degradation pathways, such as the monophenolic aldehydes produced by SCO, are expected to be toxic. Regardless of the effects on co-expression, the tight linkage of genes within MGCs may itself be driven by selection for co-adapted alleles and genes in the face of gene flow and recombination (Yeaman 2013), and the increased selectability of higher order metabolic phenotypes (Pepper 2003).

The concordant phylogenies of genes within *sco* clusters that are themselves sparsely distributed among distantly related species suggests some *sco* clusters were horizontally transferred (Reynolds *et al.* 2018) (Figure 1, Figure 3). However, the complex patterns of inheritance we observed in *sco, vao* and *gdo* gene families preclude the testing of specific HGT hypotheses (Figure 1). An alternative explanation for this pattern is the long-term retention of clusters that formed in ancient ancestors, which either were not detected in species with only a single sampled genome (because they are dispersed in the pan-genome), or were lost in most species. In either scenario, the distribution of gene clusters is a signature of selection on these specific genetic architectures. The most parsimonious explanation is that at least some horizontal *sco* cluster transfer has occurred given the large numbers of duplications and losses required to explain vertical inheritance (Szöllősi *et al.* 2015). Horizontal gene transfer (HGT) is an important source of genetic novelty for combinatorial evolution in bacteria, and is likely to impact fungal evolution as well (Soanes & Richards 2014). HGT can accelerate adaptive processes not only by giving organisms access to existing adaptive genotypes, but also by increasing combinatorial potential (Baquero 2004; Schonknecht *et al.* 2013). Clustered genes also tend to experience significantly more HGT compared with unclustered loci in fungi (Wisecaver *et al.* 2014), and HGT has been implicated in the wholesale transfer of many other fungal biosynthetic and catabolic MGCs enabling the colonization of new hosts and substrates (Slot 2017).

### Conserved specificities of alternately clustered *sco*s suggest modularity of function

It has previously been demonstrated that distantly related SCOs accommodate cleavage of similar stilbene molecules (Brefort *et al.* 2011; Díaz-Sánchez *et al.* 2013; McAndrew *et al.* 2016) (Table S9). Similarly, four distantly related *sco* homologs presented here exhibit broadly conserved activity on five of the six stilbene molecules tested (Figure 2, Table S8). Previous studies on stilbene degrading enzymes suggested that stilbene cleavage is mediated by a small number of conserved amino acid residues (Figure 2, Table S6), and the 4’-OH aromatic substitution is essential for full enzyme activity (Harrison & Bugg 2014; McAndrew *et al.* 2016). Here we show Tyr and Lys amino acid residues that mediate the hydrogen bonding with the 4’- OH, are highly conserved across fungal SCOs, with a few exceptions that merit further investigation. Further, the 4-H + 3-E residues previously shown to form the tetradentate iron coordination active site of the greater CCO family (Sui *et al.* 2015; 2016), and other amino acid residues in Tyr, Lys, and Glu containing motifs whose function is yet to be determined (Figure 2), are also highly conserved across fungal homologs. This high level of structural and functional conservation suggests that *sco* has evolved as an interchangeable metabolic module, and is consistent with the hypothesis of fungal stilbene metabolism adapting primarily through gene combination.

However, although SCOs cleave a range of 4’-OH substituted stilbenes, we show here that the breadth of substrate can vary. We observed that two closely related SCOs from *P. anserina* (model 3) and *P. roquefortii* (presumed model 3) cleave the phenylpropene isoeugenol, in addition to the conserved substrates shared with other characterized homologs. Furthermore, SCOs vary sharply in the efficiency with which they cleave piceid, a glycosylated form of resveratrol. Preservation of core functions coupled with the expansions of substrate range can minimize the change required to explore new regions of a phenotypic landscape (O’Maille *et al.* 2008; Ferrada & Wagner 2010). The recent transitions to model 3 clusters (T1-T6, AT2) might have followed relaxation of selection, which resulted in a broad-specificity SCO that was more suited to its novel role (Copley 2009)(Figure 3, Figure 4); by contrast, *Mo*SCO, which has a narrower substrate range, is anciently associated with model 2. The activity of *Pa*SCO and *Pr*SCO on isoeugenol is also interesting because the model 3 key gene, *vao*, is involved in the production of 4-hydroxycinnamyl alcohols from 4-allylphenols (e.g. eugenol and chavicol), and could indicate model 3 clusters can degrade other classes of antifungal metabolites or lignin monomers (Fraaije *et al.* 1995).

### Combinations of genes within clusters may facilitate the specialization of metabolic pathways

What could be the functional and ecological significances of the three different types of *sco* clusters? In addition to the inhibitory effects of diverse stilbenes against fungi colonizing both living and decaying plant tissues (Celimene *et al.* 1999; 2001; Jeandet *et al.* 2010), some stilbenes (e.g., piceatannol and resveratrol) were recently shown to cross-couple with monolignols during lignification of the cell wall (Del Río *et al.* 2017), where they may be expected to be released during lignin degradation processes typically carried out by saprotrophic fungi. Consequently, as particular fungal lineages frequently shift between different hosts and nutritional modes, the benefit of acquiring and optimizing mechanisms to degrade different types of stilbenes may wax and wane as populations disperse across heterogeneous environments. Transitions between a limited set of cluster types provide preliminary evidence for a constrained number of adaptive stilbene catabolism pathways, which may each represent a fitness optimum in a separate local metabolic landscape.

By decreasing recombination, metabolic gene clusters facilitate the co-adaptation between constituent genes, which may facilitate specialization on specific substrates, and possibly the formation of protein complexes that enable functional compartmentalization or metabolic channeling (Castellana *et al.* 2014). Given the evidence of evolutionary transitions between cluster types and the broad conservation of SCO activity on differently substituted stilbenes, we speculate that the specialized function of any given cluster may ultimately depend on the types of cleavage products produced by SCO’s activity on host/substrate specific stilbenes. Predicted functions encoded in each *sco* cluster suggest accessory genes enable degradation of alternatively substituted monophenolic cleavage products by alternative, conserved ring-cleavage pathways (Figure S15). Model 2 cluster gene functions (*gdo, fph*) suggest a plausible pathway for transformation to citric acid cycle intermediates by gentisic acid cleavage (Greene *et al.* 2014). Model 1 and model 3 cluster gene functions (e.g. *dhbd* and *vao,* respectively) suggest ring cleavage of downstream products by the β-ketoadipate pathway, which also leads to the production of citric acid cycle intermediates (Mäkelä *et al.* 2015). Specialization of *sco* clusters to particular cleavage products may circumvent fitness defects arising from cross-talk with other (especially core) metabolic pathways (McGary *et al.* 2013; Keller 2015). Cluster distributions in large pan-genomes, dispersed among large populations and through horizontal transfer is expected to increase the strength of ecological selection that results in this specialization and optimization (Lynch 2006). At the same time, these large gene pools would provide broad access to combinatorial partners that are selected under similar environmental conditions (Baquero 2004).

Together, our results are consistent with a process where combinatorial evolution through genomic rearrangements facilitates fungal adaptation to plant defense chemistry, and highlight the additional importance of enzymes with modular functions. Future work to characterize these stilbene catabolic pathways in full, and to assess their contributions to fitness in particular environments, will help to refine our understanding of the processes contributing to metabolic adaptation and specialization.

## ACKNOWLEDGEMENTS

We thank the staff of the Molecular and Cellular Imaging Center at The Ohio State University. Special thanks to Mike Kelly (MCIC, Columbus) for expert advice and help with UPLC measurements. We thank Marysabel Mendez Acevedo for her help in cloning the *Podospora anserina* SCO. We thank Thomas K. Mitchell, Enrico Bonello, Ana Paula Alonso, and Laura Kubatko for their kind support in providing materials and helpful comments for parts of this work. We thank Cathie Aime, Daniele Armaleo, Jose Maria Barrasa, Stephen B. Goodwin, Gerald Bills, Burt H. Bluhm, Gregory Bonito, Sarah Branco, Tom Bruns, Kathryn Bushley, Laurie Connell, Pedro Crous, Gunther Doehlemann, Paul Dyer, David Ezra, Dave Greenshields, Igor Grigoriev, Alexey Grum-Grzhimaylo, Andrii Gryganskyi, Colleen Hansel, David S. Hibbett, Patrik Inderbitzin, Bjorn D. Lindahl, Daniel L. Lindner, Francois Lutzoni, Jon Karl Magnuson, Francis Michel Martin, Don Natvig, Kerry O’Donnell, Robin Ohm, Amy Jo Powell, Marie-Noëlle Rosso, Trey Sato, Steven W. Singer, Joseph Spatafora, John W. Taylor, Adrian Tsang, Gillian Turgeon, Rytas Vilgalys, and Kenneth H. Wolfe for providing access to genomic data prior to publication; these sequence data were produced in collaboration with the user community by the U.S. Department of Energy Joint Genome Institute, a DOE Office of Science User Facility, supported by the Office of Science of the U.S. Department of Energy under Contract No. DE-AC02-05CH11231. Computational work was performed using the resources of the Ohio Supercomputer Center. This work was supported by the Fonds de Recherche du Quebec-Nature et Technologies (EGT), the NSF (DEB-1638999) and an OARDC SEEDS Interdisciplinary Team Research Competition Grant.

